# Identification and characterization of a temperature sensitive chlorotic soybean mutant

**DOI:** 10.1101/2024.02.02.578604

**Authors:** C. Nathan Hancock, Tetandianocee Germany, Priscilla Redd, Jack Timmons, Jeffery Lipford, Samantha Burns, Sergio Alan Cervantes-Perez, Marc Libault, Wenhao Shen, Yong-qiang Charles An, Lisa Kanizay, Melinda Yerka, Wayne A. Parrott

## Abstract

Screening a transposon-mutagenized soybean population led to the discovery of a recessively inherited chlorotic phenotype. This “vir1” phenotype results in smaller stature, weaker stems, and a smaller root system with smaller nodules. Genome sequencing identified 15 candidate genes with mutations likely to result in a loss of function. Amplicon sequencing of a segregating population was then used to narrow the list to a single candidate mutation, a single-base change in *Glyma.07G102300* that disrupts splicing of the second intron. Single cell transcriptomic profiling indicates that this gene is expressed primarily in mesophyll cells and RNA sequencing data indicates it is upregulated in germinating seedlings by cold stress. Previous studies have shown that mutations to *Os05g34040*, the rice homolog of *Glyma.07G102300*, produced a chlorotic phenotype that was more pronounced in cool temperatures. Growing soybean vir1 mutants at lower temperatures also resulted in a more severe phenotype. In addition, transgenic expression of wild type *Glyma.07G102300* in the knockout mutant of the Arabidopsis homolog *At4930720* rescues the chlorotic phenotype, further supporting the hypothesis that the mutation in *Glyma.07G102300* is causal of the vir1 phenotype.

## Introduction

Pressure to increase agriculture yields is increasing as human populations grow, even as arable land and fertilizer resources remain limited. The genetic improvement of crop species to overcome these challenges will be facilitated by the identification of the genes controlling crop yield, quality, and stress tolerance (Thomson, Ismail et al. 2010). This is a challenging task because plant genomes contain thousands of genes, many of which are essential and cannot be deleted, or are functionally redundant with other genes. Identifying gene function using traditional genetic mapping has been a slow and labor-intensive process, requiring thousands of plants to generate the recombination events needed to pinpoint the gene(s) of interest. Thus, faster methods for identifying the genes responsible for altered or mutant phenotypes are needed. Here we describe an example in which advances in genomic sequencing and analysis facilitated the identification of a gene responsible for a chlorotic mutant soybean. Uncovering the pathways responsible for healthy chloroplast production has the potential to increase photosynthetic capacity and unlock additional yield potential.

Soybeans are a source of both oil and protein, making them valuable for the food supply, animal production, and renewable fuels. Scientists are developing soybean gene discovery resources to identify targets for breeding and genome editing. These include mutagenic populations made with traditional chemical [i.e. EMS or NMU (Cooper, Till et al. 2008, Espina, Ahmed et al. 2018)] and radiation [i.e. fast neutron (Bolon, Haun et al. 2011)] mutagens as well as the development of transposon tagging resources based on the *mPing, Tnt1*, and *Ac/Ds* transposable elements (Mathieu, Winters et al. 2009, Hancock, Zhang et al. 2011, Cui, Barampuram et al. 2013, Johnson, McAssey et al. 2021). In addition, site directed mutagenesis techniques that can target specific sequences have been developed for soybean (Jacobs, LaFayette et al. 2015, Du, Zeng et al. 2016, Slotkin, Liu et al. 2023). As all these methods result in off-target mutations, it is important to develop an efficient pipeline for whole genome sequence analysis to complement these studies so that plants having the fewest off-target mutations may be selected for downstream analysis and breeding.

Virescent mutants, in which young leaves fail to develop the normal green color, are an easily observable phenotype. They have been identified in rice (Sugimoto, Kusumi et al. 2004, Yoo, Cho et al. 2009, Gong, Su et al. 2014, Sun, Zheng et al. 2017), tobacco (Archer and Bonnett 1987), cotton (Benedict, McCree et al. 1972), maize (Chollet and Paolillo Jr 1972) cucumber (Miao, Zhang et al. 2016), Arabidopsis (Brusslan and Tobin 1995, Koussevitzky, Stanne et al. 2007, Zhou, Cheng et al. 2009), and peanut (Alberte, Hesketh et al. 1976). Evaluation of the underlying genes causing virescent phenotypes commonly shows that these genes are generally associated with chloroplast development. The ability of plants to efficiently synthesize and maintain chlorophyll is an important indicator of plant health and is one of the most common phenotypes collected by agronomists and growers to monitor the irrigation and fertilization status of crops. Thus, identifying the genes important for soybean chloroplast development is economically significant.

This report describes the discovery and genomic analysis of a virescent soybean mutant, vir1, discovered in a mutagenic population. This is an example of how the advances in sequencing technology allowed for the gene underlying the mutant phenotype to be identified in a relatively small plant population. The description of these techniques provides a potential template to help plant scientists solve the complex problem of sorting through genome sequencing data to identify specific mutations. The causal gene identified from this analysis can be used to further understand the requirements for healthy chloroplast development.

## Materials and Methods

### Plant Transformation and Screening

Transformation of “Jack” (Nickell, Noel et al. 1990) soybeans with *mPing*-based mutagenesis plasmids was performed as previously described (Hancock, Zhang et al. 2011). The resulting plants were screened for phenotypes in research plots at the University of Georgia for multiple years under standard growing conditions, which includes at least 6 weeks of supplemental lighting to prevent early flowering. In 2019, chlorotic plants (Shot 125-94A-3-27-2-7-2) were crossed as the male to control Jack plants. The resulting F_2_ seedlings (Cross 19-1) were phenotyped in the greenhouse before transplanting into the field. A chlorotic F_2_ plant was crossed as a male to Williams 82 to produce the Cross 20-7 F_2_ population. F_3_ progeny from green seedlings were grown to identify lines segregating for the vir1 phenotype.

### Genome Sequencing and Bioinformatics

Genomic DNA was purified from immature leaves using a scaled up version of the CTAB method as described previously (Johnson, McAssey et al. 2021). The RNA and other contaminants were removed by a second purification with the E.Z.N.A. Plant DNA DS Mini Kit (Omega Bio-Tek). The sequencing libraries were prepared using Illumina TruSeq DNA PCR-Free LP kits and were sequenced on an Illumina HiSeq.

The sequence analysis pipeline was based on https://gencore.bio.nyu.edu/variant-calling-pipeline-gatk4/. Briefly, the raw reads were assembled to the Williams 82 reference genome [Gmax_275_v2.0.fa (Schmutz, Cannon et al. 2010)], duplicates were marked and sorted with Picard, and variant SNPs and indels were identified with GATK. BCFtools was used to compare the variants found in the different genomes. SnpEff was used to identify SNPs and INDELs that are likely to cause a loss in protein function. Annotated scripts are available at: https://github.com/cnhancock/Plant-Genome-Resequencing/ The raw sequence reads are available at: https://submit.ncbi.nlm.nih.gov/subs/sra/SUB13607166/overview

### Amplicon Sequencing

DNA from six vir1 progeny plants from Cross 20-7 and Cross 20-8 (Williams 82 x vir1) was pooled and amplified with primers that flank known polymorphisms in the candidate genes and contain the Illumina Adapter (IA) sequence (Supplemental Table 1). The resulting fragments were purified and sent for Illumina sequencing.

### Genotyping

Purified genomic DNA was used for PCR with Chr07:9757891 For and Chr07:9758683 Rev primers (Supplemental Table 1). The resulting PCR products were digested with *Dra*I before agarose gel electrophoresis.

### Single Cell Expression Analysis

The average expression of *Glyma.07G102300* was extracted from all cell clusters from the tissues composing the soybean single-cell atlas (unpublished data; Libault lab) by importing the RDS files using Seurat (Hao, Hao et al. 2021). The results were visualized as a transcriptomic profile using a heatmap plot from yellow (low expression) to red (high expression) across the entire soybean single-cell atlas.

### RNA Analysis

RNA was purified from leaf tissue from plants at the V3 stage. The tissue (0.1g) was ground in 800 µl of Trizol using ZR Bashing Bead Lysis Tubes (Zymo Research). The Direct-zol RNA Miniprep Plus Kit (Zymo Research) was then used for RNA isolation according to the manufacturer’s recommendations (Zymo Cat# R2070S). cDNA was generated from 1 μg of RNA using the iScript Reverse Transcription Supermix for RT-qPCR kit (BioRad). PCR with Chr07:9757891 For and Chr07 9758683 Rev primers was performed using APEX master mix. The resulting bands were gel purified and cloned into pGEM T-easy (Promega) and sent out for Sanger sequencing.

### Arabidopsis Overexpression

The PCR fragment produced by high fidelity PCR of wild type soybean with the Glyma.07G102300 Rev attb and Glyma.07G102300 For attb primers (Supplemental Table 1) was cloned into pDONR Zeo, sequenced, then transferred into a modified version of the pEarleyGate 100 plasmid (Earley, Haag et al. 2006) with the 35S promoter replaced with the RSP5a promoter (pEG 100R) (Renken, Mendoza et al. 2023). This construct was transformed into GV3031 Agrobacteria by triparental mating with a pRK2013 helper strain and selecting on YM supplemented with Rifampicin and Kanamycin.

The Arabidopsis T-DNA mutant CS883546 was genotyped by performing PCR (Supplemental Figure 1) with the At4G30720 For and At4G30720 Rev primers (Supplemental Table 1). Seeds from a line identified as homozygous null for *At4G30720* were transformed using the floral dip method (Clough and Bent 1998).

### Image Analysis

LED strip lighting was added to incubators to create identical growth chambers at different temperatures. Soybean seeds were inoculated with commercial inoculant, then planted 3 cm deep in Sungrow Professional Growing Mix with no additional fertilizer. The first fully opened trifoliate leaf from each plant was digitally imaged from 23 cm above the leaf. The images were cropped to only include the region with the leaf, and processed through Batch Load Image Processor (BLIP) software (Renfroe-Becton, Kirk et al. 2022) to identify and characterize leaf pixels. Statistical analysis was performed with JMP statistical software.

### Transcriptome Analysis

Transcriptome sequencing data were downloaded from the National Center for Biotechnology Information (NCBI) database and were analyzed as previously described (Goettel, Zhang et al. 2022). The transcript accumulation of each gene was normalized and indicated in transcript per million (TPM) based on the *G. max* reference genome [Wm82.a2.v1 (Schmutz, Cannon et al. 2010)]. Linear modeling was used to analyze the expression data from Bioproject PRJNA999924. For the other datasets, a two-tailed T-test was performed to identify treatments with > 2-fold change in average transcript accumulation and p-value ≤ 0.05 in comparison with their controls.

### Supplemental Data files

Supplemental Table 1 – Table of primer sequences used in this study

Supplemental Figure 1 – PCR analysis of CS883546 Arabidopsis

Supplemental Figure 2 – Changes in nodulation

## Results

### Identification of the vir1 Mutant

A chlorotic mutant was discovered in a soybean population (Jack cultivar) transformed with an *mPing*-based activation tag and the ORF1 and Transposase genes required for its mobilization (Hancock, Zhang et al. 2011, Johnson, McAssey et al. 2021). This “vir1” phenotype was especially apparent in young plants (early in the growing season) and when grown in the greenhouse [during the winter, inoculated, no additional fertilizer] (Figure 1A). In addition to chlorotic leaves, the vir1 plants had a smaller root system (Figure 1B) and fewer nodules that appeared to be smaller (Supplemental Figure 2). The observed changes in root architecture suggested that nutrient uptake or nitrogen fixation may be responsible for chlorosis. However, leaf nutrient analysis, supplemental fertilization, and grafting experiments did not produce any clear results. Analysis of 47 seedlings from a segregating F_3_ population showed that 29% were chlorotic, consistent with a 3:1 ratio caused by a single recessive allele (χ^2^ = 0.574, *p* = 0.449). When grown in field conditions, the vir1 plants were significantly shorter at maturity (Figure 1C), and more plants were observed to lodge due to broken stems. Together, these observations are consistent with the vir1 plants having decreased overall photosynthetic capacity.

**Figure 1.**
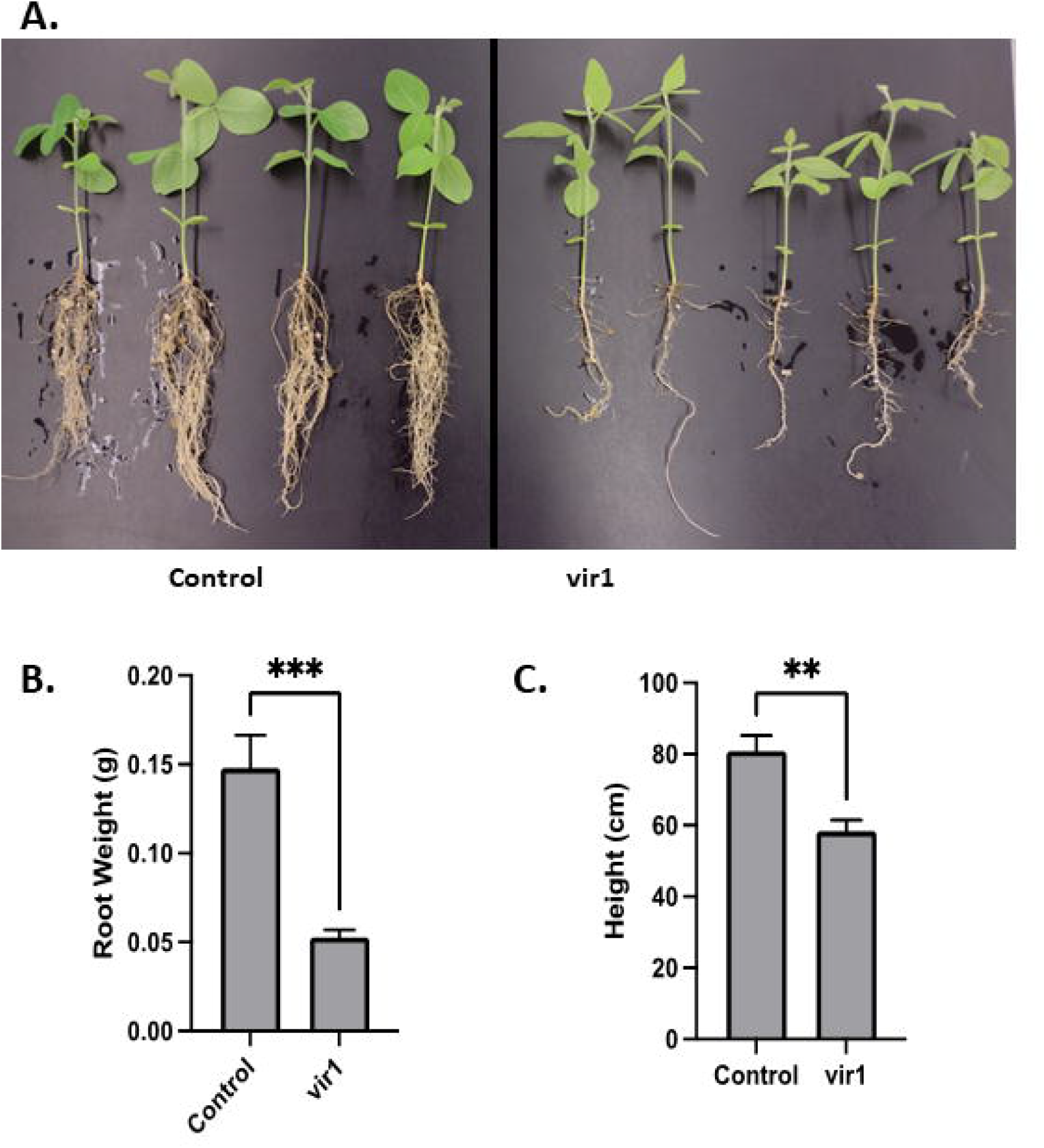
Phenotype of vir1 soybeans. A: Images of greenhouse grown Control (left) and vir1 (right) seedlings. B: Average root weight for the greenhouse grown plants shown in A. C: Average height of mature field grown plants. Error bars represent the standard error of the mean. Asterisks indicate a significant difference based on a two tailed t-test.

PCR analysis of these small segregating populations indicated that the vir1 phenotype was not associated with the transgenes present in the population (i.e., they potentially resulted from random, *de novo* mutation). To identify the causal gene, we sequenced the genome of a vir1 plant and assembled it to the Williams 82 reference genome (Schmutz, Cannon et al. 2010). The variants present in the vir1 genome were identified using a modified version of a previously developed bioinformatics pipeline (Khalfan 2020). Using sequences from two other Jack samples, we identified the SNPs and indels that were unique to the vir1 plant. Using SnpEff, we identified 15 variants that were predicted to disrupt gene function (Table 1). Five of these putative knockout variants were in genes that had previously been reported in the *Glycine max* haplotype map list of genes with known loss of function alleles (Torkamaneh, Laroche et al. 2021). Because the cultivars used for the haplotype map were homozygous and not chlorotic, this indicated that these five variants were unlikely to be causal for the vir1 phenotype. To narrow the list further, we designed primers flanking the 10 remaining candidate variants that would amplify a 300 bp fragments including the variant locus. We then amplified these fragments from a pool of six F_3_ plants exhibiting the vir1 phenotype (a bulk segregant analysis) and sequenced the amplicons. Analysis of these sequences indicated that the variant co-segregating with the vir1 phenotypes was a single base polymorphism in *Glyma.07G102300* (Table 1). The finding that the mutant allele for *Glyma.07G102300* was only present in 86.6% of the reads from the pool suggests that one of the plants in the pool was misidentified as chlorotic. This misidentification was likely the effect of other environmental factors, such as fertility or disease, that also contribute to chlorophyll production.

**Table 1.** Variants present in *vir1*.

| Chromosome Position | Reference Sequence | Variant Sequence | Haplotype Map LOF | % Mutant in Pool | Gene |
| --- | --- | --- | --- | --- | --- |
| Chr02:15034955 | G | A |  | 20 | *Glyma.02G145600* |
| Chr03:45596156 | TG | T | Yes | -- | *Glyma.03G262600* |
| Chr06:15970458 | C | A |  | 44.6 | *Glyma.06G184400* |
| Chr06:37844984 | C | A | Yes | -- | *Glyma.06G234900* |
| Chr07:9758094 | C | T |  | 86.6 | *Glyma.07G102300* |
| Chr07:39528933 | A | T |  | 18 | *Glyma.07G220700* |
| Chr08:3701874 | C | T |  | 78 | *Glyma.08G047300* |
| Chr09:3336685 | G | A |  | 40.9 | *Glyma.09G040000* |
| Chr10:51324749 | G | A | Yes | -- | *Glyma.10G296400* |
| Chr12:35671633 | ATTG | A | Yes | -- | *Glyma.12G195200* |
| Chr14:44340827 | G | A |  | 21 | *Glyma.14G180600* |
| Chr18:4953932 | C | T |  | 10.4 | *Glyma.18G056600* |
| Chr18:12258084 | C | T |  | 2 | *Glyma.18G107900* |
| Chr19:47236307 | G | A | Yes | -- | *Glyma.19G220100* |
| Chr19:48663487 | G | A |  | 5.7 | *Glyma.19G237800* |

The mutation in the mutant allele of *Glyma.07G102300* contains a *Dra*I site that is absent from the wild type allele. We developed a cleaved amplified polymorphic sequence (CAPS) marker by using PCR and digestion with *Dra*I enzyme to genotype plants from an F_3_ population segregating for vir1 (Figure 2). Results confirmed that the chlorotic phenotype is observed in plants that are homozygous for the variant but not in plants that are heterozygous for the mutation. This result suggests that the homozygous recessive Glyma.07G102300 allele is 100% correlated with the vir1 phenotype. Consequently, the CAPS marker is a perfect (functional) co-dominant marker for the vir1 trait.

**Figure 2.**
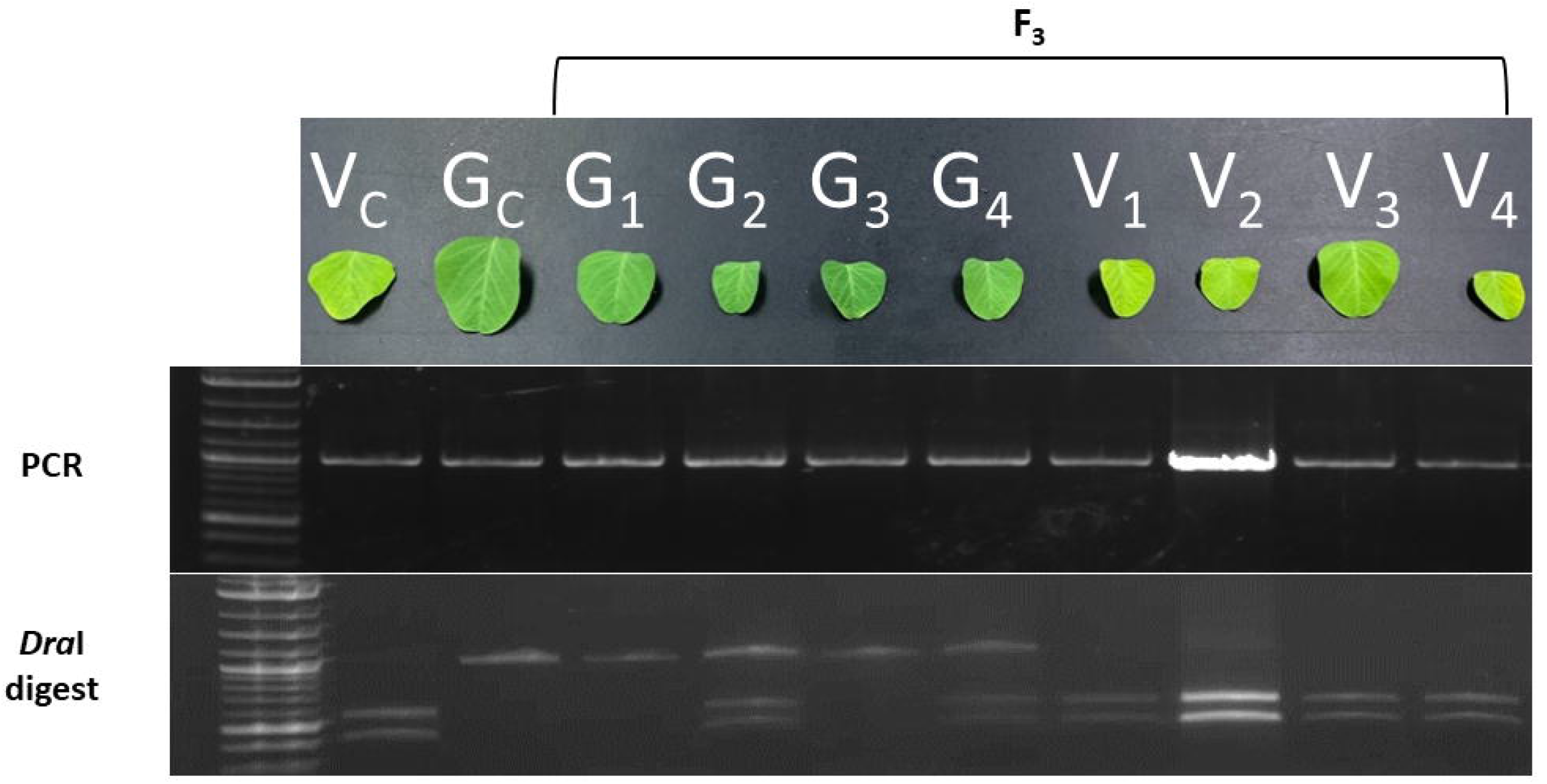
Association of the *Glyma.07G102300* variant with the vir1 phenotype. PCR and *Dra*I digestion analysis of *Glyma.07G102300* from an F_3_ population segregating for the vir1 phenotype. V = vir1, G = green, vir1 control (Vc) = original vir1 mutant, green control (Gc) = Jack.

### *Glyma.07G102300* mRNA analysis

The protein coding sequence of *Glyma.07G102300* is highly conserved across a range of plant species (Figure 3A), suggesting that it plays an essential role in plant biology. Soybean is an ancient tetraploid, so it was not surprising to find that *Glyma.07G102300* has a closely conserved paralog, *Glyma.04G155100*. However, the predicted coding of the chromosome 4 sequence is missing a large portion of the N-terminal domain due to the presence of a large transposable element insertion (Figure 3A). This likely makes the *Glyma.04G155100* version nonfunctional, suggesting that *Glyma.07G102300* is the only functional copy in the soybean genome.

**Figure 3.**
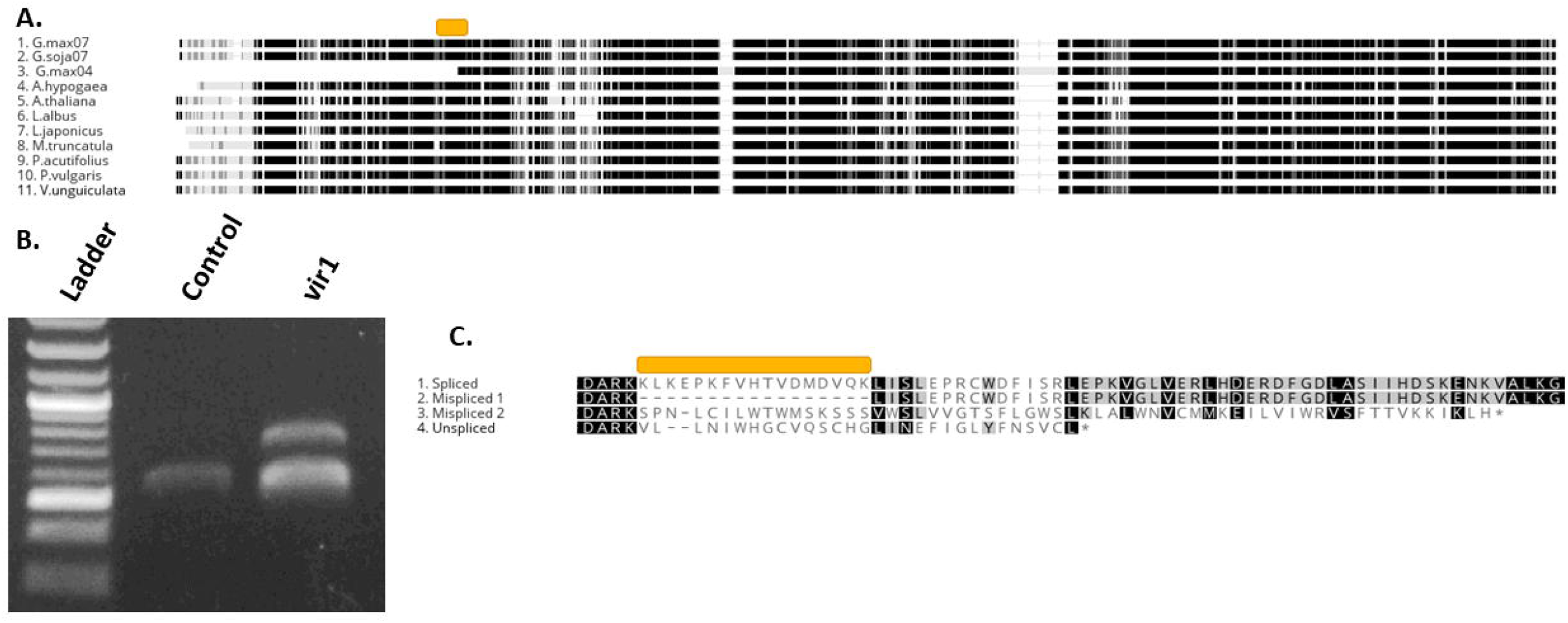
*Glyma.07G102300* mRNA analysis. A: Protein alignment of *Glyma.07G102300* homologs from various dicot species. Black regions are conserved, orange box indicates region missing from the misspliced 1 mRNA. B: Gel of rtPCR amplicons from *Glyma.07G102300*. Ladder = 1 Kb+ (New England Biolabs). C: Protein alignment of the translated cDNAs cloned from vir1 shown in B.

The knockout allele of *Glyma.07G102300* described herein contains a single C to T change at position 9758094 (Table 1) that is predicted to disrupt the second intron splice acceptor site. We compared the splicing of *Glyma.07G102300* in wild-type and vir1 mutants by isolating RNA, making cDNA, and performing PCR with primers that flank the second intron (Figure 3B). Our results show that the wild-type allele of *Glyma.07G102300* mRNA is spliced as expected, giving a single band. However, the vir1 allele mRNA resulted in two bands. Sequencing the two bands showed that the upper band was an RNA with an unspliced intron 2 that results in a frame shift and early termination (Figure 3C). Sequencing the lower bands showed that they were aberrantly spliced in two ways, one version resulting in the loss of 17 conserved amino acids and the other producing a frame shift and early stop (Figure 3C). This result is consistent with the hypothesis that vir1 is caused by disrupting the function of *Glyma.07G102300*.

To further elucidate the function of *Glyma.07G102300*, we took advantage of the soybean single cell atlas (unpublished data) to determine the expression levels across the cell clusters. Figure 4 shows a heatmap indicating expression levels across a wide range of cell types. The highest expression of wild-type *Glyma.07G102300* is found in four clusters composing the mesophyll cells in trifoliate leaves. This is consistent with our phenotypic observations in leaves, and also points to a potential role in chloroplasts which are abundant in these tissues. This also matches prior proteome analysis results showing that the wild-type Arabidopsis homolog, *At4g30720*, was present in chloroplast stroma (Zybailov, Rutschow et al. 2008). Expression of an N-terminal GFP fusion to *At4930720* was also reported in the chloroplast (Vlad, Rappaport et al. 2010). The rice homolog, *Os05g34040*, has a chloroplast targeting signal and both C-terminal GFP and N-terminal CFP fusions were localized to chloroplasts (Morita, Nakagawa et al. 2017, Sun, Zheng et al. 2017). Together, these results suggest that Glyma.07G102300 protein is likely localized to the chloroplast.

**Figure 4.**
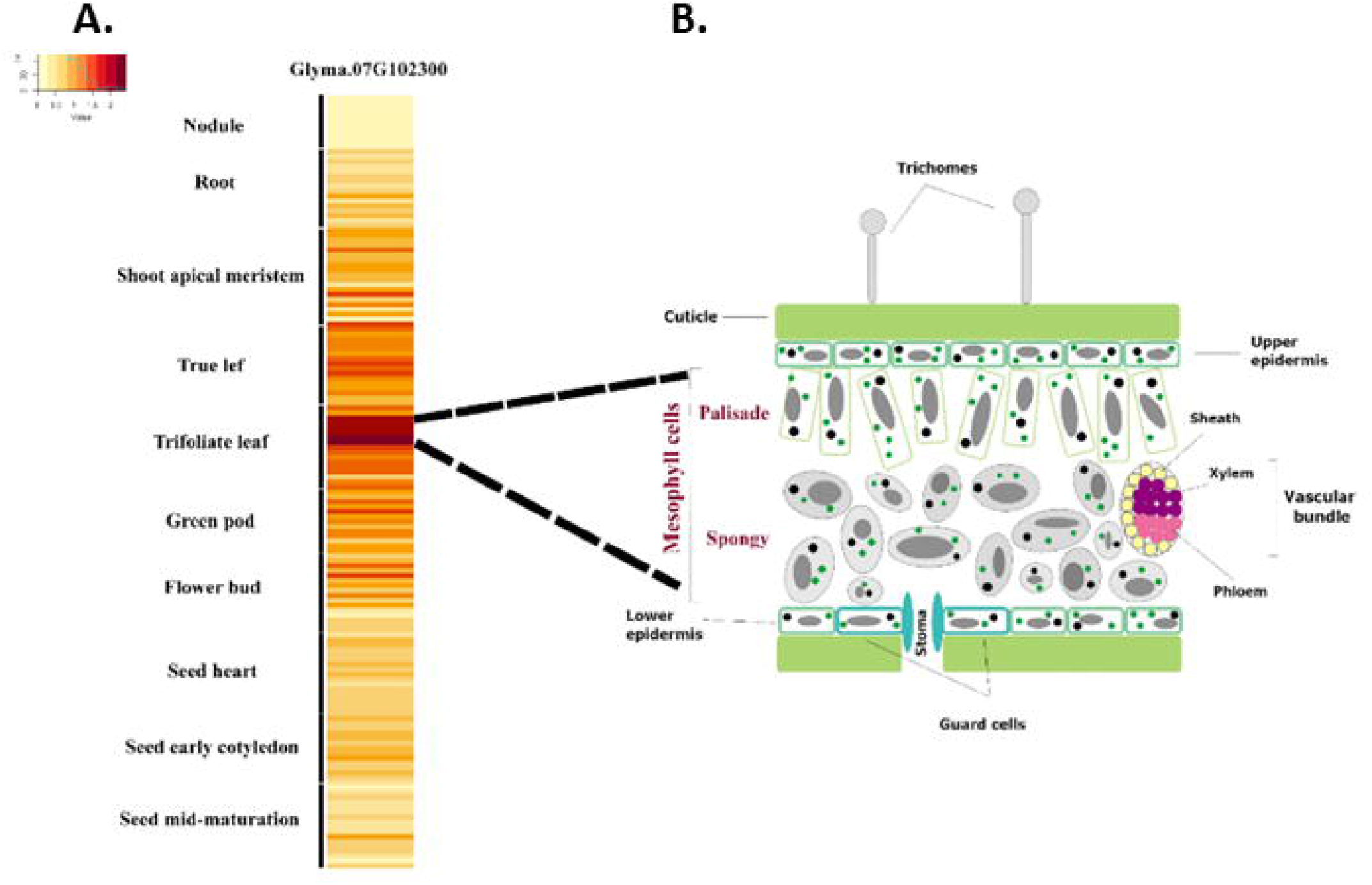
Single cell expression analysis of *Glyma.07G102300* in soybean. A: Heatmap of the average expression of *Glyma.07G102300* in each soybean cell type (y-axis). B: Diagram representing the cellular diversity of the soybean trifoliate leaf. The palisade and spongy mesophyll cells both show high *Glyma.07G102300* expression activity.

### Functional Analysis

The protein produced by *Glyma.07G102300* is predicted to be an oxidoreductase or electron carrier-like protein. Previous studies have identified chlorotic mutants caused by disruption of *Glyma.07G102300* homologs. The Arabidopsis homolog, *At4930720*, was identified as a pigment defective (pde327-1) SALK line [SALK_059716, CS883546 (Meinke 2020)]. Variation in *At4930720* between Col-0 and Bur-0 ecotypes was found to be the underlying cause of a chlorotic and 70% smaller phenotype observed in recombinant inbred lines generated from those parents (Vlad, Rappaport et al. 2010). This study showed that some ecotypes contain two functional copies of *At4930720*, but at least one functional copy is required for normal pigmentation and growth (Vlad, Rappaport et al. 2010). To test if the soybean and Arabidopsis versions of these genes were functionally the same (protein sequence identity = 66.8%, similarity = 87.6%) we cloned and overexpressed a wild-type genomic *Glyma.07G102300* gene in CS883546, the knockout mutant of *At4930720* (Figure 5). We observed an increase in plant size in the hemizygous T_1_ generation and used the T_2_ seeds segregating for a single copy of the transgene to quantitate the increase. Figure 5 shows representative plants with and without the transgene. The number of pixels per plant (background removed) was significantly higher when the *RPS5a:Glyma.07G102300* transgene was present. Similar results were obtained when the rice homolog, *Os05g34040*, was overexpressed in SALK_059716, the parental line for CS883546 (Wang, Zhang et al. 2016), indicating it is functionally conserved across dicot and monocot plant species.

**Figure 5.**
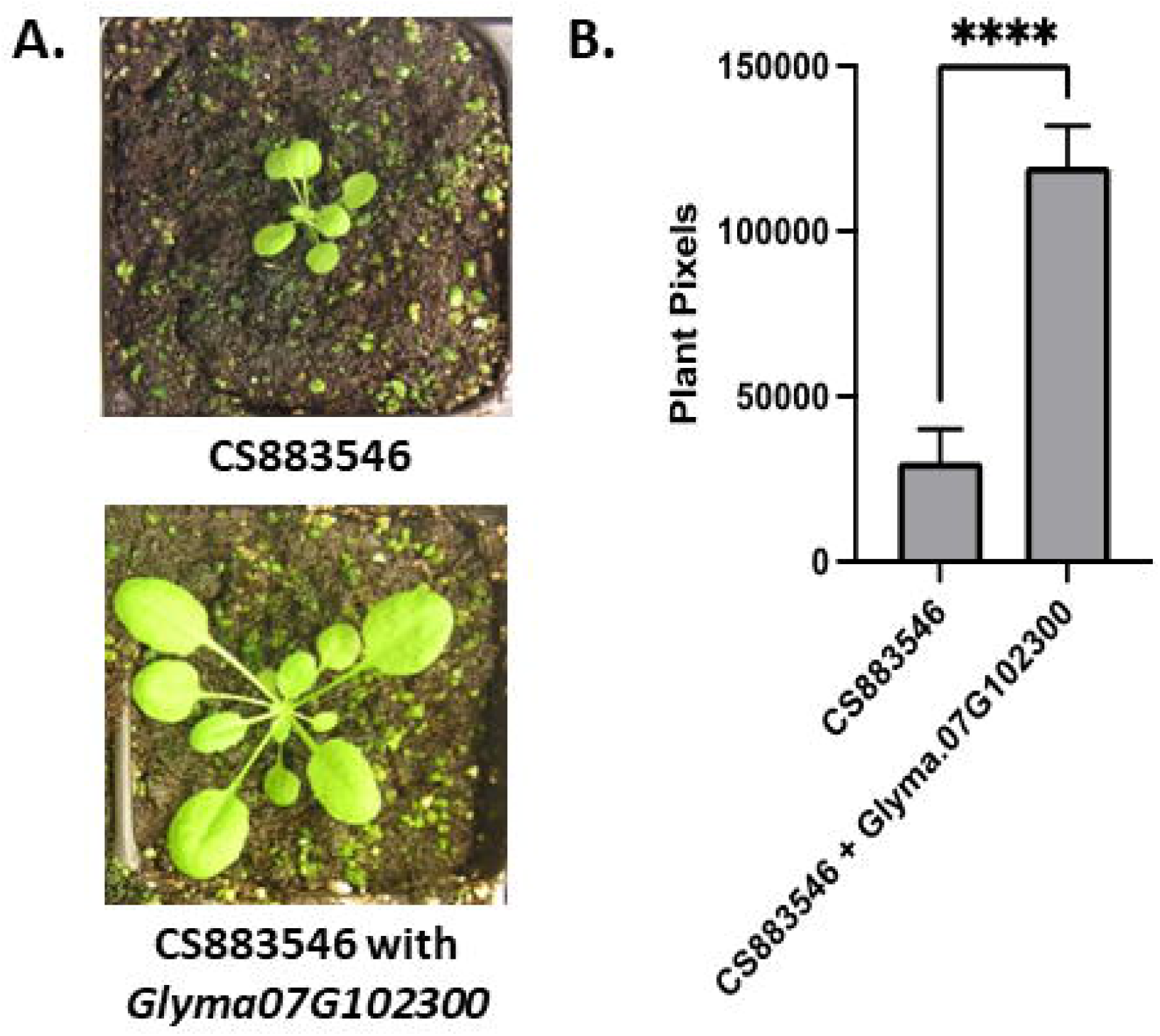
Rescue of the *At4g30720* loss of function mutant with a *Glyma.07G102300* transgene. A: Images of the CS883546, the *At4g30720* loss of function mutant, with (bottom) and without (top) the *RPS5A:Glyma.07G102300* expression construct. B: Average plant size of eight seedlings of each type measured by the number of plant pixels present in the images. Error bars represent the standard error of the mean. Asterisks indicate a significant difference based on a two tailed t-test.

Multiple groups have reported that the chlorotic phenotype from loss of function mutations in *Os5G34040* were more pronounced in cool temperatures [20°C] than in warmer temperatures [30°C] (Wang, Zhang et al. 2016, Morita, Nakagawa et al. 2017, Sun, Zheng et al. 2017). Based on this information, we tested if temperature influenced the phenotype of the soybean vir1 mutants (Figure 6). We observed small differences in leaf color and shape between the control and vir1 mutants at 25°C and 30°C. However, when we grew the mutant at 18°C, we observed a drastic difference in leaf color, shape, and area (Figure 6). This temperature sensitivity further supports the hypothesis that the mutation we identified in *Glyma.07G102300* is causal of the vir1 phenotype.

**Figure 6.**
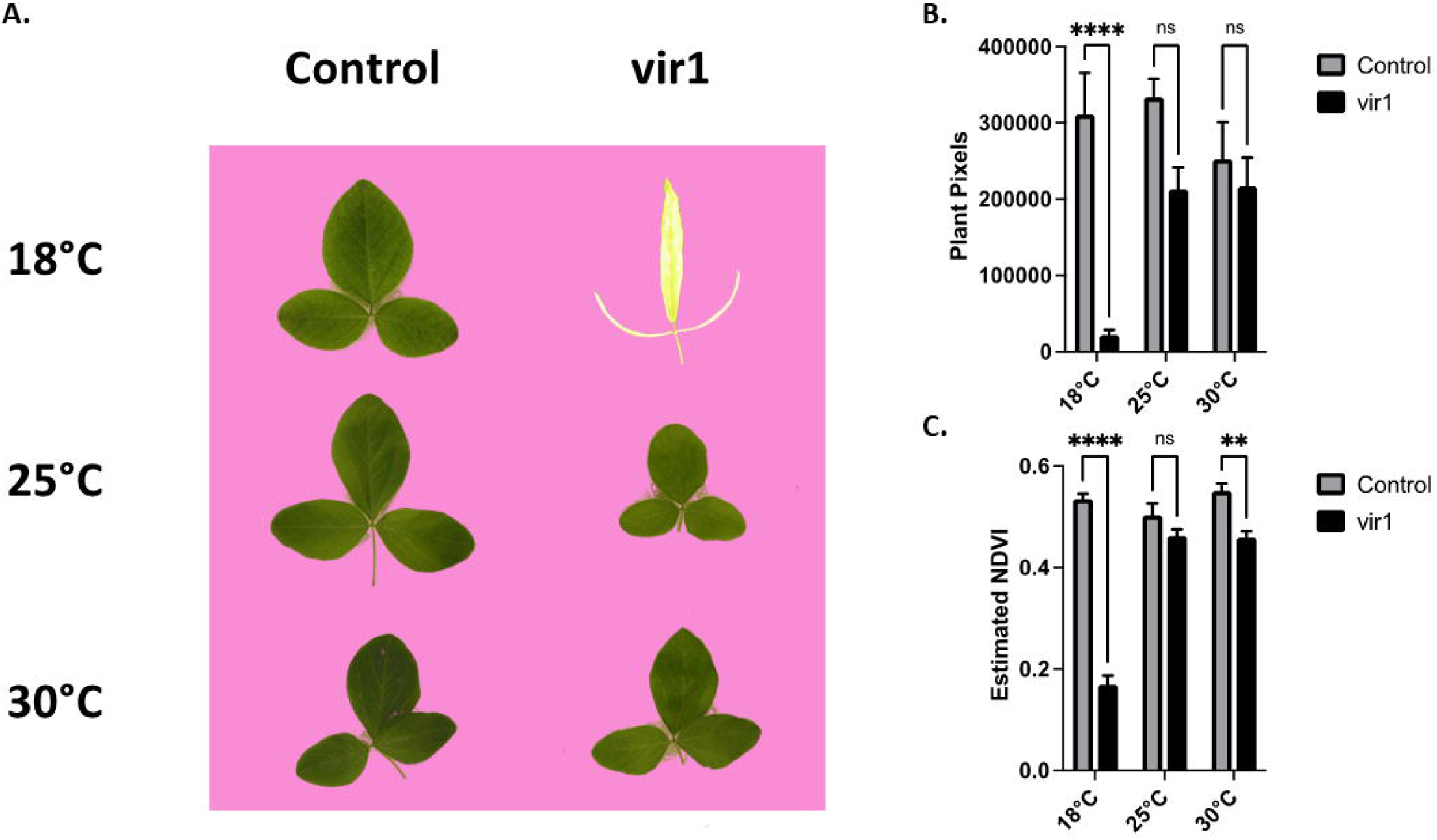
Temperature effect on leaf size and color. A: Example images of first trifoliate leaves grown at three temperatures with identical lighting. Background pixels have been replaced with pink to indicate the plant pixels used for image analysis. B: Leaf size measured as the average number of plant pixels. C: Average predicted leaf Normalized Difference Vegetation Index (NDVI) based on the color of the plant pixels. Averages represent 5 or 6 trifoliate leaves. Error bars represent the standard error of the mean. Asterisks indicates a significant difference based on a two-way ANOVA with Šídák’s multiple comparisons test, ns = not significantly different.

Previous reports on the expression of the rice homolog *Os05g34040*, indicate that lower temperatures increased expression (Sun, Zheng et al. 2017), but a similar study observed no change in expression (Morita, Nakagawa et al. 2017). To gain additional insight into the expression pattern of *Glyma.07G102300*, we analyzed leaf RNA expression data from publicly available data sets. The most relevant data set we identified was Bioproject # PRJNA999924 that includes replicated control (20°C) and low temperature (6°C) treatments of germinating seeds [two genotypes (Zheng, Xie et al. 2023)] over 48 hours (Figure 7). Linear modeling of *Glyma.07G102300* expression from this dataset showed that there was a significant interaction between time and treatment (t = 4.687, *p*<0.001), so the control and low temperature treatments were modelled separately. The slope of transcripts per million (TPM) vs. time for all of the control data showed no significant difference from zero (t = 7.364, *p*<0.001). However, we detected a significant positive relationship between TPM vs. time for the low treatment (t= 9.021, *p*<0.001). Thus, this analysis suggests that *Glyma.07G102300* expression is induced by cold treatment in germinating soybean seeds. In addition to this result, the public data indicated other treatments that significantly alter *Glyma.07G102300* expression (Table 2). A leaf development experiment using the Hedou12 cultivar (Feng, Yang et al. 2021) showed that unexpanded trifoliate leaves had less *Glyma.07G102300* expression than fully expanded trifoliate leaves. This corelates with the normal greening observed during leaf development. In addition, multiple experiments showed a decrease in *Glyma.07G102300* expression under drought stress (Table 2), consistent with lower photosynthetic activity and growth. Similarly, an RNA expression dataset comparing mock and Soybean Mosaic Virus inoculated leaves (Chen, Arsovski et al. 2016) indicates that viral infection results in a drop in *Glyma.07G102300* expression. Surprisingly, an increase in *Glyma.07G102300* expression was observed 120 hours after inoculation with *P. pachyrhizi*, the fungal agent responsible for Asian soybean rust.

**Table 2.**
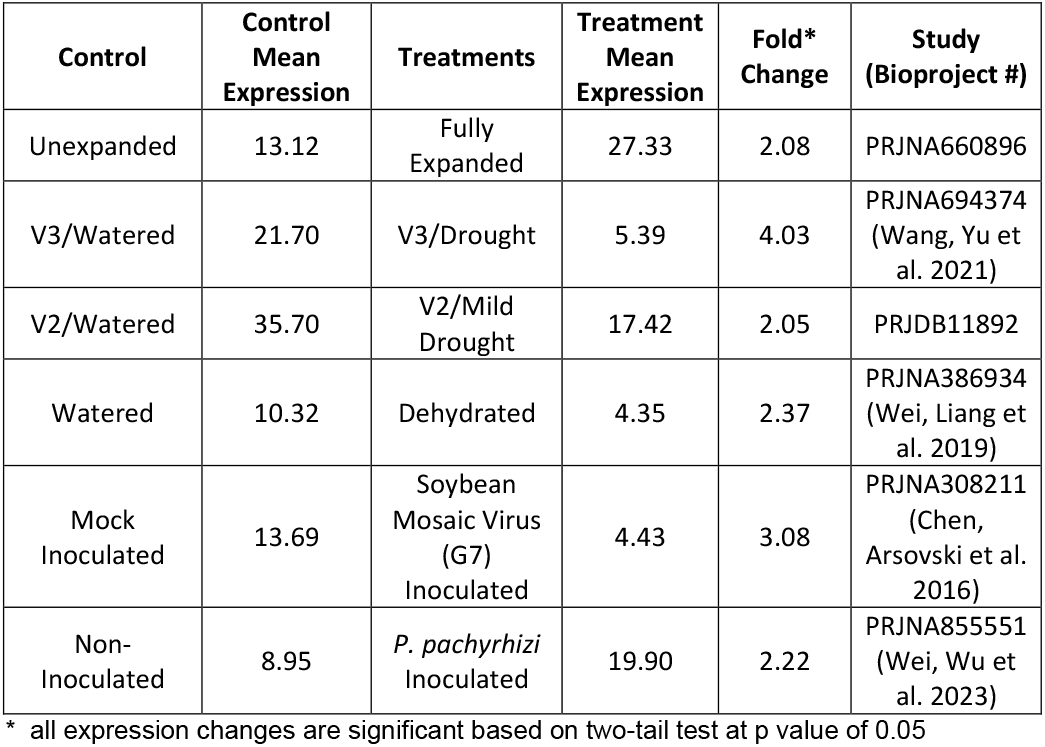
Gene expression changes for *Glyma.07G102300* in trifoliate leaves.

| Control | Control Mean Expression | Treatments | Treatment Mean Expression | Fold* Change | Study (Bioproject #) |
| --- | --- | --- | --- | --- | --- |
| Unexpanded | 13.12 | Fully Expanded | 27.33 | 2.08 | PRJNA660896 |
| V3/Watered | 21.70 | V3/Drought | 5.39 | 4.03 | PRJNA694374 (Wang, Yu et al. 2021) |
| V2/Watered | 35.70 | V2/Mild Drought | 17.42 | 2.05 | PRJDB11892 |
| Watered | 10.32 | Dehydrated | 4.35 | 2.37 | PRJNA386934 (Wei, Liang et al. 2019) |
| Mock Inoculated | 13.69 | Soybean Mosaic Virus (G7) Inoculated | 4.43 | 3.08 | PRJNA308211 (Chen, Arsovski et al. 2016) |
| Non-Inoculated | 8.95 | *P. pachyrhizi* Inoculated | 19.90 | 2.22 | PRJNA855551 (Wei, Wu et al. 2023) |

\* all expression changes are significant based on two-tail test at p value of 0.05

**Figure 7.**
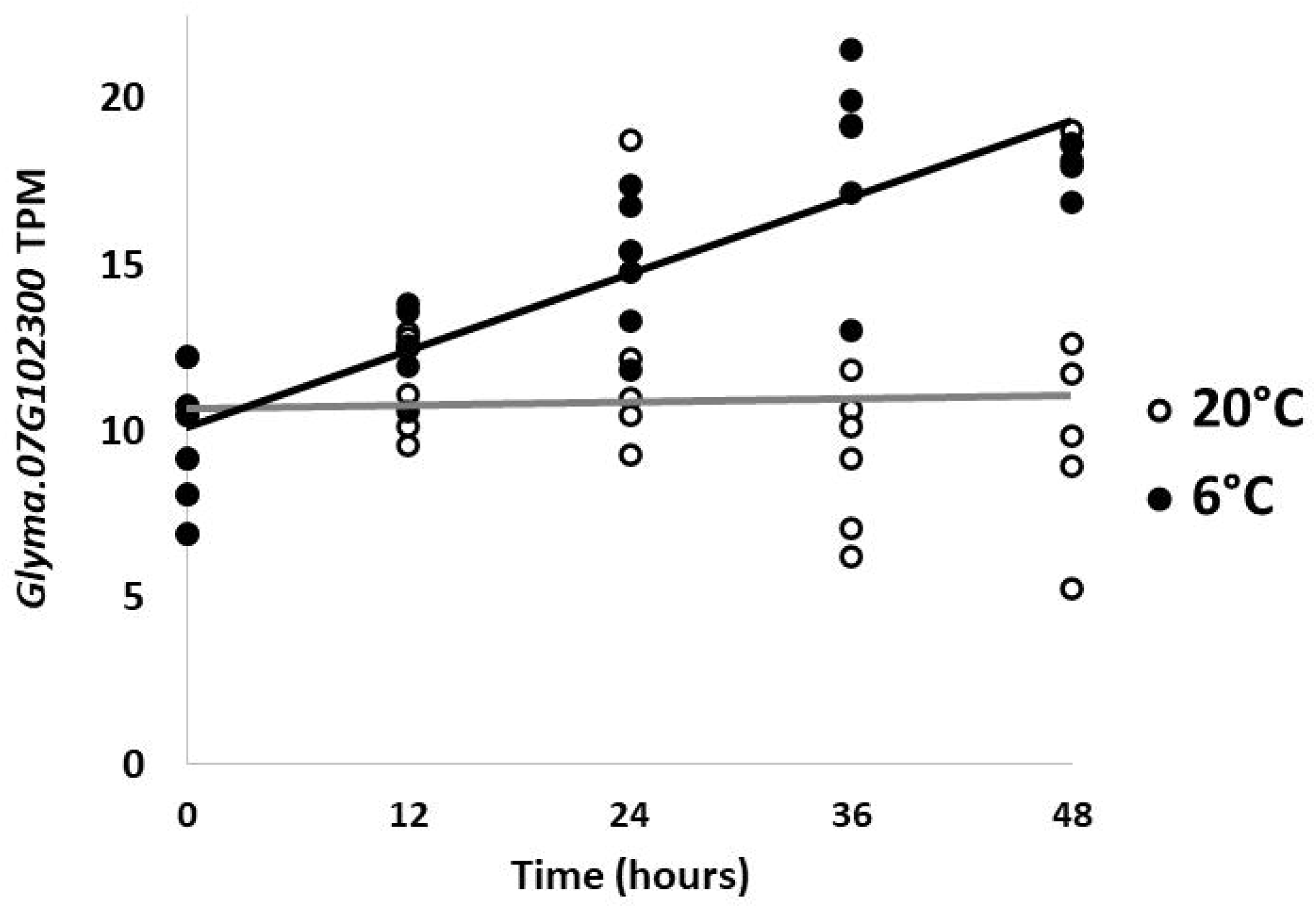
Expression of *Glyma.07G102300* under cold stress. Plot of *Glyma.07G102300* expression in germinating seeds measured in transcripts per million (TPM) from Bioproject #PRJNA999924 under control (20°C) and low temperature (6°C) conditions. Each time point includes 6 data points [three each from chromosome fragment substitution line R48 and R89 (Zheng, Xie et al. 2023)]. The linear regression for 20° and 6°C are represented by a gray and black line, respectively.

Additional expression studies, including cold stress treatment, will be helpful in elucidating the role of the *Glyma.07G102300*.

## Discussion

Our results provide a successful example of using whole genome resequencing followed by bulk segregant analysis of targeted amplicons to identify the causal mutation for an economically impactful plant mutant phenotype. This technique should be applicable to a wide range of plant species that have an available reference genome. This project required whole genome sequencing a single vir1 mutant plant and comparing it to a pool of wild-type plants with the same genetic background (Jack). The need for wild-type whole genome sequences would have been reduced if the mutation had occurred in the reference Willaims 82 cultivar. However, detailed characterization of Williams 82 plants has also shown that intercultivar genetic heterogeneity is pervasive (Haun, Hyten et al. 2011), which could complicate candidate gene discovery.

One factor that facilitated this study was the ease with which the vir1 phenotype could be scored. Even with this benefit, our amplicon sequencing suggests that we mischaracterized one of the plants included in our pool. The phenotypic selection was performed with plants grown at about 25°C, where the phenotype is less severe. We now know that this mistake could have been avoided by growing the plants at even cooler temperatures (i.e., 18°C). Despite this mistake, our methods clearly identified the causal mutation. This suggests that application of this technique to other less obvious phenotypes should be feasible if bulk segregant analysis is based on amplicon sequencing of pools of extreme phenotypes. We acknowledge that a single-gene trait in a diploid species (lacking a functional copy from its ancient genome duplication) is the easiest scenario for developing and sequencing extreme phenotypic pools. In the case of more quantitative traits, or in polyploid species, a similar approach would still be effective at capturing large-effect loci to quickly narrow down the list of candidate genes [Reviewed in (Wang, Han et al. 2023)].

Taken together, our results suggest that the knockout allele of *Glyma.07G102300* is the cause of the vir1 phenotype. Prior soybean mutant studies identified a number of chlorophyl deficient mutants named y1 through y20 (Palmer, Pfeiffer et al. 2004). The y8 and y10 phenotypes were both reported to be more severe in young plants and segregate in a 3:1 ratio (Woodworth and Williams 1938, Probst 1950), consistent with the vir1 phenotype. Unfortunately, no mapping data is available for the y8 and y10 mutants, and they are no longer available to the soybean research community due to the lack of a soybean stock center. Thus, to our knowledge, vir1 is the only virescent soybean mutant with a known locus. However, mapping studies The lack of other virescent mutants in soybean may be due to the gene redundancy present in this ancient tetraploid. In fact, the vir1 mutant would probably have not been observed if the homologous *Glyma.04G155100* gene was not altered (Figure 3A). The fact that soybean and its wild ancestor *Glycine soja* only have one functional copy of this conserved oxidoreductase gene could be due to random loss, or it could be due to selection for moderate expression. We did not detect any adverse effects from overexpression of the soybean *Glyma.07G102300* gene in Arabidopsis (Figure 5), and some Arabidopsis accessions naturally have two copies of the *At4g30720* homolog (Vlad, Rappaport et al. 2010). Our analysis of publicly available data indicates that cold stress upregulates *Glyma.07G102300* expression in germinating seedlings (Figure 7). However, additional experiments are needed to clarify if expression also increases in fully expanded leaves that already have higher levels of *Glyma.07G102300* transcript (Table1).

The dramatic expression of the vir1 phenotype under cool temperatures (Figure 6) suggests that the gene is critical to chloroplast development under cold stress. One possible explanation is that this oxidoreductase plays a role in protecting the chloroplasts from reactive oxygen species (ROS). Photosynthesis produces additional ROS under cold stress conditions, resulting in oxidative damage (Tambussi, Bartoli et al. 2004, Banerjee and Roychoudhury 2019). The loss of oxidoreductase protein may exacerbate ROS stress to the point of inhibiting chloroplast development. Another possibility is that *Glyma.07G102300* is important for the expression of essential chloroplast genes. Previous study of the rice homolog *Os05g34040* indicated that plastid-encoded RNA polymerase activity is decreased in the knockout plants (Sun, Zheng et al. 2017). However, it is not clear if this is causal, or simply a downstream effect of chloroplast stress.

To our knowledge, this is the first study to report decreased root biomass (Figure 1) and decreased nodulation (Supplemental Figure 2) associated with a virescent mutant in soybean. We hypothesize that the reduced root biomass could exacerbate the leaf phenotype by reducing nutrient uptake from the soil. Nitrogen deficiency is known to cause leaf chlorosis, so any reduction in nodulation is likely to exacerbate chlorosis. While our results point to a role for *Glyma.07G102300* in the leaf, additional studies on the function of this important oxidoreductase protein’s role in roots are warranted.

## Acknowledgements

Special thanks to Michael Shtutman and Hao “Emily” Ji at the USC Functional Genomics Core for assistance with genomic library preparation, sequencing, and bioinformatics. We are grateful to Alec Sherratt (University of South Carolina Aiken) and Kendall Kirk (Clemson University) for helping with image processing and statistical analysis. We thank Rick Meyer for technical assistance in analysis of the RNA-seq data and Derek Zelmer (University of South Carolina Aiken) for assistance with linear modeling. This project was supported by grant P20GM103499 (SC INBRE) from the National Institute of General Medical Sciences, National Institutes of Health. It was also facilitated by award #1444581 from the National Science Foundation.

