## Supplemental Table 1 for "Identification and characterization of a temperature sensitive chlorotic soybean mutant"

| Variant | Gene | Forward Primer | Reverse Primer |
|---|---|---|---|
| Chr02:15034955 | Glyma.02G145600 | ACACTCTTTCCCTACACGACGCTCTTCCGATCTGATGAGAACTCGAAGAAAAGCAAG | GACTGGAGTTCAGACGTGTGCTCTTCCGATCTATGAGGCAAGAGAAATCTCATTCA |
| Chr06:15970458 | Glyma.06G184400 | ACACTCTTTCCCTACACGACGCTCTTCCGATCTTCCACCAAAGTTGAATTTCTCAAG | GACTGGAGTTCAGACGTGTGCTCTTCCGATCTATTGAAGCAAAATCATAGAGCCAC |
| Chr07:9758094 | Glyma.07G102300 | ACACTCTTTCCCTACACGACGCTCTTCCGATCTTCCCCTTTCAATGCAACTTTATTT | GACTGGAGTTCAGACGTGTGCTCTTCCGATCTAACAGTAAACAAAATGCGGCTAAT |
| Chr07:39528933 | Glyma.07G220700 | ACACTCTTTCCCTACACGACGCTCTTCCGATCTTGGCATATTCAGGAGACATGTAAC | GACTGGAGTTCAGACGTGTGCTCTTCCGATCTTGATGAACCCAAAAATTGCTGATT |
| Chr08:3701874 | Glyma.08G047300 | ACACTCTTTCCCTACACGACGCTCTTCCGATCTATAGAAAATCCCTCCCAAACCTTT | GACTGGAGTTCAGACGTGTGCTCTTCCGATCTGATTCATATCCGACTCCTGATTGA |
| Chr09:3336685 | Glyma.09G040000 | ACACTCTTTCCCTACACGACGCTCTTCCGATCTGTCAAGAAGAAGGTTATTGTGGTG | GACTGGAGTTCAGACGTGTGCTCTTCCGATCTTCTGGTGGATTTGTTCAAGGTTAT |
| Chr14:44340827 | Glyma.14G180600 | ACACTCTTTCCCTACACGACGCTCTTCCGATCTCATGTCCGGGGATAAGATAGAAAA | GACTGGAGTTCAGACGTGTGCTCTTCCGATCTATTCAGCACAATGAAAAACACATG |
| Chr18:4953932 | Glyma.18G056600 | ACACTCTTTCCCTACACGACGCTCTTCCGATCTGAGTGTTACACGAGTGATGATCTA | GACTGGAGTTCAGACGTGTGCTCTTCCGATCTGAAACTTGAGAAGCTCTTGTCTTC |
| Chr18:12258084 | Glyma.18G107900 | ACACTCTTTCCCTACACGACGCTCTTCCGATCTTGGGTGATTTTATCATCATGCAAC | GACTGGAGTTCAGACGTGTGCTCTTCCGATCTCCAAAAACCACATTGAGATTGTGA |
| Chr19:48663487 | Glyma.19G237800 | ACACTCTTTCCCTACACGACGCTCTTCCGATCTTTTCAAATTGGCTGAATCCCAAAA | GACTGGAGTTCAGACGTGTGCTCTTCCGATCTTTTGCAAGTTCCTGTTGAATCTTT |
| | Glyma.07G102300 attB | GGGGACAAGTTTGTACAAAAAAGCAGGCTTCATGTTCCTTCTTCCTTCTCCTTCGTCC | GGGGACCACTTTGTACAAGAAAGCTGGGTCCTAGTACTTGACAACTCCAACATTTTGAGC |
| | Chr07 9757691-9758683 | ACACTCTTTCCCTACACGACGCTCTTCCGATCTTCCCCTTTCAATGCAACTTTATTT | CCTTATGGTAGTGGCAAGCTAATA |
| | At4G30720 | GACGCTCCTT GAGTTAGAGC CTCG | CCCCAAAGCG AACCAACGTT TTCA |
| | AtChr3:20585893 - 20586281 | TATGCAGCCACGTCAATGTCCCAT | CGTCAGATAATTAGTTGGTTGAGCCCT |
