## Supplemental Figures for "Identification and characterization of a temperature sensitive chlorotic soybean mutant"

**_Supplemental Figure 1._ PCR analysis of CS883546 Arabidopsis**

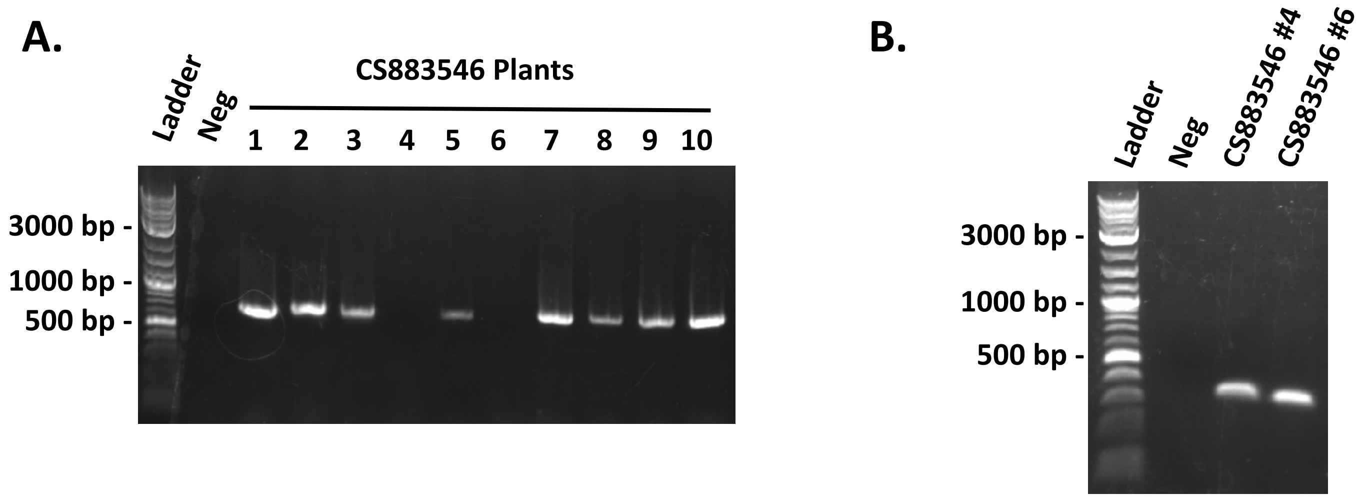


Gel electrophoresis images of PCR identification CS883546 plants that were homozygous for T-DNA insertion in the *At4G30720* gene. A: Ten CS883546 plants screened with the At4G30720 For and At4G30720 Rev primers which amplifies the wild type At4G30720 gene. B: Confirmation of the presence of DNA for CS883546 samples #4 and #6 in part A by using AtChr3:20585893 For and AtChr3:20586281 Rev primers. Ladder = NEB 1kb+ Quickload DNA Ladder, Neg = Water Control.

*Supplemental Figure 2.* **Changes in nodulation**

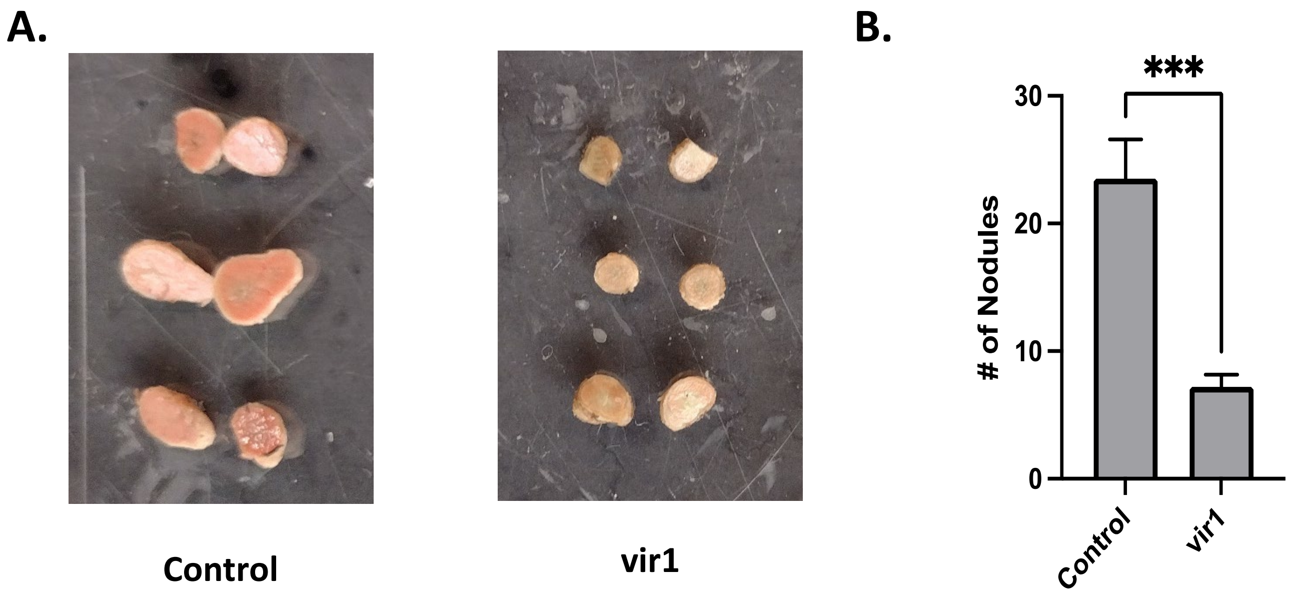


A:  Images of halved nodules from control and vir1 seedlings from Figure 1. B: Average number of nodules observed on the roots of the plants in Figure 1.  Error bars represent the standard error of the mean. Asterisks indicates a significant difference based on a two tailed t-test.
